# A population of adult satellite-like cells in *Drosophila* is maintained through a switch in RNA-isoforms

**DOI:** 10.1101/174151

**Authors:** Hadi Boukhatmi, Sarah Bray

## Abstract

Adult stem cells are important for tissue maintenance and repair. One key question is how such cells are specified and then protected from differentiation for a prolonged period. Investigating the maintenance of *Drosophila* muscle progenitors (MPs) we demonstrate that it involves a switch in *zfh1/ZEB1* RNA-isoforms. Differentiation into functional muscles is accompanied by expression of *miR*-*8/miR*-*200*, which targets the major *zfh1*-*long* RNA isoform and decreases Zfh1 protein. Through activity of the Notch pathway, a subset of MPs produce an alternate *zfh1*-*short* isoform, which lacks the *miR*-*8* seed site. Zfh1 protein is thus maintained in these cells, enabling them to escape differentiation and persist as MPs in the adult. There, like mammalian satellite cells, they contribute to muscle homeostasis. Such preferential regulation of a specific RNA isoform, with differential sensitivity to miRs, is a powerful mechanism for maintaining a population of poised progenitors and may be of widespread significance.

## INTRODUCTION

Growth and regeneration of adult tissues depends on stem cells, which remain undifferentiated while retaining the potential to generate differentiated progeny. For example, muscle satellite cells (SCs) are a self-renewing population that provides the myogenic cells responsible for postnatal muscle growth and muscle repair (Chang and Rudnicki, 2014). One key question is how tissue specific stem cells, such as satellite cells, are able to escape from differentiation and remain undifferentiated during development, to retain their stem cell programme though-out the lifetime of the animal.

It has been argued that the progenitors of *Drosophila* adult muscles share similarities with satellite cells and thus provide a valuable model to investigate mechanisms that maintain stem cell capabilities (Aradhya, et al., 2015; Figeac, et al., 2007). After their specification during embryogenesis, these muscle progenitors (MPs) remain undifferentiated throughout larval life before differentiating during pupal stages. For example, one population of MPs is associated with the wing imaginal disc, which acts as a transient niche, and will ultimately contribute to the adult flight muscles. These MPs initially divide symmetrically to amplify the population. They then enter an asymmetric division mode in which they self-renew and generate large numbers of myoblasts that go on to form the adult muscles (Gunage, et al., 2014). In common with vertebrates, activity of Notch pathway is important to maintain the MPs in an undifferentiated state (Gunage, et al., 2014; Mourikis and Tajbakhsh, 2014; Mourikis, et al., 2012; Bernard, et al., 2010). To subsequently trigger the muscle differentiation program, levels of Myocyte Enhancer factor 2 (Mef2) are increased and Notch signalling is terminated (Elgar, et al., 2008; Bernard, et al., 2006). Until now it was thought that all MPs followed the same fate, differentiating into functional muscles. However, it now appears that a subset persist into adulthood forming a population satellite-like cells ((Chaturvedi, et al., 2017) and see below). This implies a mechanism that enables these cells to escape from differentiation, so that they retain their progenitor-cell properties.

The *Drosophila* homologue of ZEB1/ZEB2, ZFH1 (zinc-finger homeodoman 1), is a candidate for regulating the MPs because this family of transcription factors is known to repress Mef2, to counteract the myogenic programme (Postigo and Dean, 1999; Postigo, et al., 1999). Furthermore, *zfh1* is expressed in the MPs when they are specified in the embryo and was shown to be up-regulated by Notch activity in an MP-like cell line (DmD8) (Figeac, et al., 2010; Krejci, et al., 2009). Furthermore, an important regulatory link has been established whereby microRNAs (miRs) are responsible for down-regulating ZEB/Zfh1 protein expression to promote differentiation or prevent metastasis in certain contexts (Zaravinos, 2015; Vandewalle, et al., 2009). For example, the miR-200 family is significantly up-regulated during type II cell differentiation in fetal lungs, where it antagonizes ZEB1 (Benlhabib, et al., 2015). Likewise, *miR*-*8*, a *miR*-*200* relative, promotes timely terminal differentiation in progeny of Drosophila intestinal stem cells by antagonizing *zfh1* and *escargot* (Antonello, et al., 2015). Conversely, down-regulation of *miR*-*200* drives epithelial mesenchymal transition (EMT) to promote metastasis in multiple epithelial derived tumours (Korpal, et al., 2008; Park, et al., 2008). Such observations have led to the proposal that the ZEB/miR-200 regulatory loop may be important in the maintenance of stemness, although examples are primarily limited to cancer contexts and others argue that the primary role is in regulating EMT (Antonello, et al., 2015; Brabletz and Brabletz, 2010). The MPs are thus an interesting system to investigate whether this regulatory loop is a gatekeeper for the stem cell commitment to differentiation.

To investigate the concept that ZEB1/Zfh1 could be important in sustaining progenitor-type status, we examined the role and regulation of *zfh1* in *Drosophila* MPs/SCs. Our results show that *zfh1* plays a central role in the maintenance of undifferentiated MPs and, importantly, that is expression is sustained in a population of progenitors that persist in adults (pMPs). Specifically these pMPs express an alternate short RNA isoform of *zfh1* that cannot be targeted by *miR*-*8*. In contrast, the majority of larval precursors express a long isoform of *zfh1*, which is subject to regulation by *miR*-*8* so that Zfh-1 protein levels are suppressed to enable differentiation of myocytes. Expression of alternate *zfh1*-*short* isoform is thus a critical part of the regulatory switch to maintain a pool of progenitor “satellite-like” cells in the adult. This type of regulatory logic, utilizing RNA isoforms with differential sensitivity to miRs, may be of widespread relevance for adult stem cell maintenance in other tissues.

## RESULTS

### Zfh1 is required for maintenance of muscle progenitors

*As zfh1* was previously shown to antagonize myogenesis it was a plausible candidate to maintain the muscle progenitor cells in *Drosophila* and prevent their differentiation. We therefore examined its expression in the muscle progenitors (MPs) associated with the wing disc. Zfh1 was clearly present throughout the large group of MPs, which can be distinguished by the expression of Cut (Ct) (Figure 1A-A”). At early stages Zfh1 is uniformly expressed throughout the MPs (Figure supplement 1) but at later stages the levels become reduced in the cells with high Cut expression (Figure 1 A”). These cells give rise to the direct flight muscles (DFMs), whereas the remaining MPs, where Zfh1 expression is high, give rise to the indirect flight muscles (IFMs) (Figure 1 A”; (Sudarsan, et al., 2001)). Zfh1 expression in MPs is therefore regulated in a manner that correlates with different differentiation programmes.

**Figure 1.**
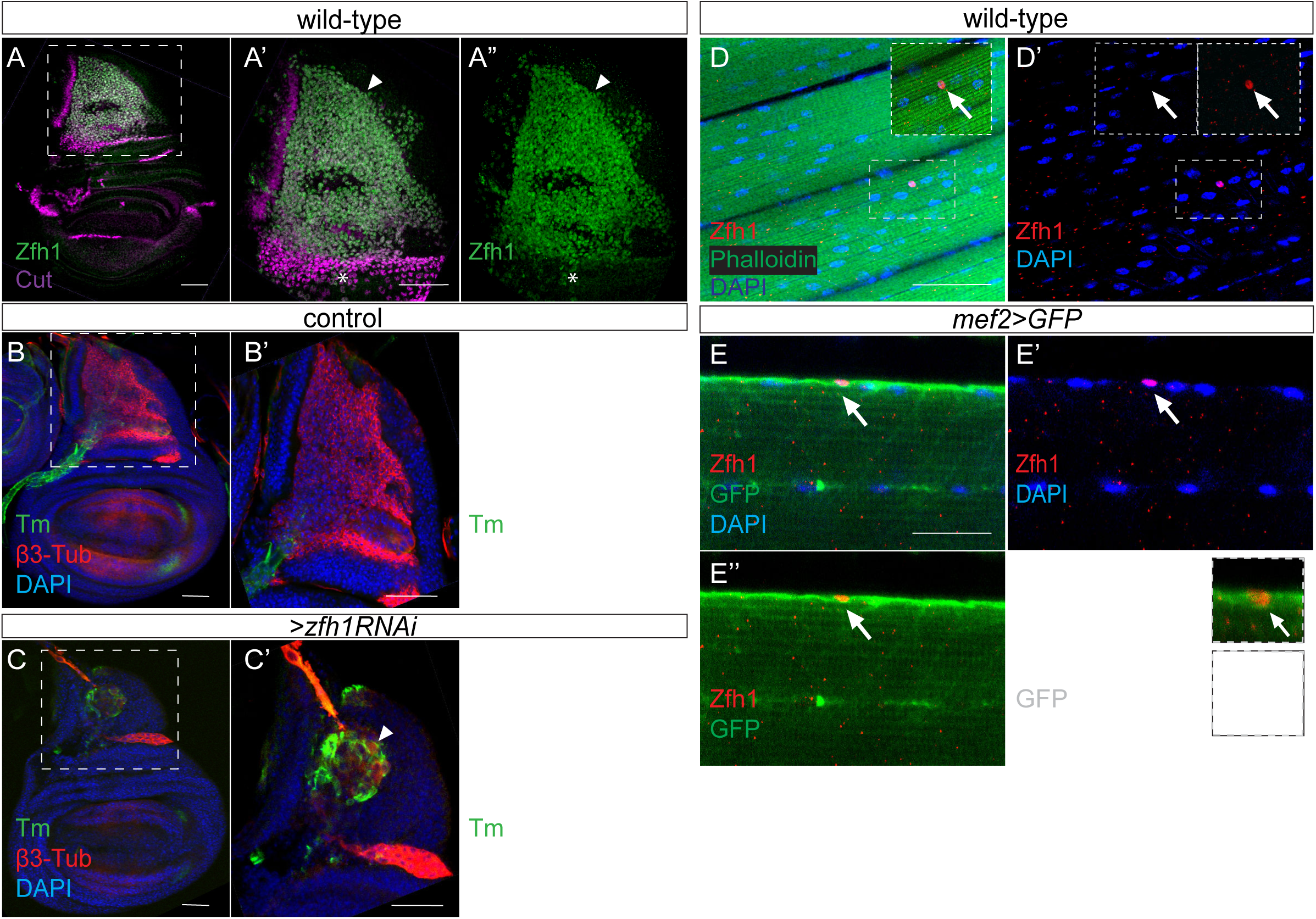
Zfh1 expression and function in MPs and in adult pMPs. **(A-A”).** Zfh1 (Green) and Cut (Purple) expression in MPs associated with third instar wing discs, **(A’-A”).** higher magnification (3X) of boxed region in A. Zfh1 is present in all MPs, but those with highest Cut expression have lower levels of Zfh1 (asterisk). Scale bars: 50 μM. **(B-C).** Down regulation of *zfh1* in MPs induces premature differentiation of the MPs (arrowhead in C’-C”). β3-Tubulin (β3-Tub, red) and Tropomyosin (Tm, Green) expression in control (B, *1151*-*Gal4*>*UAS*-*wRNAi*) and Zfh1 depleted (C, *1151*-*Gal4*>*UAS*-*Zfh1RNAi*) third instar wing discs, (B’-C”) higher magnification (3X) of boxed regions in B and C. **(D-D’).** Zfh1 expression (red) indicates the existence of persistent muscle progenitors (pMPs; arrows) associated with the muscle fibers (Phalloidin (Green), DNA/Nuclei (Blue)). The immune cell marker P1 was included in the immunostaining and is absent from the pMPs (Figure supplement 1). Scale bars: 50 μM. **(E-E”’).** Zfh1 (Red) expressing pMPs (e.g. arrows in E”’) are closely embedded in the muscle lamina of the adult indirect flight muscles and express Mef2 (myogenic cells; *Mef2*-*Gal4*>*UAS*-*Src::GFP*, green). Nuclei (Blue), Scale bars: 25 μM.

To determine whether Zfh1 is required in the MPs to antagonize myogenic differentiation we tested the consequences from expressing a hairpin RNA to target/silence *zfh1 specifically* in MPs (using *1151*-*Gal4* as a driver). Tropomyosin (Tm), a protein normally expressed in differentiated muscles, was clearly detectable when *zfh1*-*was* down-regulated (Figure 1B-C). A similar result was obtained using a Myosin Heavy Chain (MHC) reporter, which revealed that small muscle fibers formed precociously in ~ 20% of *zfh*-*1* depleted wing discs (n=4/20 wing discs; Figure supplement 1). Consistent with the premature expression of muscle differentiation markers, β-3Tubulin staining showed that decreased *zfh1* led to abnormal MP cell morphology (Figure 1B’-C’). These results demonstrate that reduced *zfh1* expression causes MPs to initiate the muscle differentiation program indicating that *zfh1* is required to prevent MP differentiation.

Lineage tracing experiments suggest that a subset of wing disc MPs have characteristics of muscle stem cells and remain undifferentiated even in adult *Drosophila* (Chaturvedi, et al., 2017; Gunage, et al., 2014). Since Zfh-1 is necessary to prevent differentiation in MPs, we reasoned that its expression might be sustained in these persistent muscle progenitors (pMPs). Indeed, some individual cells, closely associated with adult IFM muscle fibers, had high levels of Zfh1 expression whereas the differentiated muscle nuclei exhibited no detectable expression (Figure 1D-D’). To better characterise these Zfh1-positive adult cells, we expressed a membrane-tagged GFP (*UAS*-*Src::GFP*) under the control of a specific muscle driver *mef2*-*Gal4* (*mef2*>*GFP*). This confirmed that Zfh1 was expressed in *mef2*>*GFP* expressing cells, indicating they were myogenic, and that these cells were closely embedded in the muscle lamina (Figure 1E-E”’). These cells therefore have characteristics of persistent muscle progenitors and likely correspond to the so-called adult satellite cells recently identified by others (Figure 1D and E). Although many of the Zfh1 expressing cells were clearly co-expressing Mef2>GFP, Zfh1 was also detected in another population that lacked Mef2 expression. Often clustered, these cells were co-labelled with a plasmatocyte marker P1/Nimrod indicating that they are phagocytic immune cells (Figure supplement 2).

The results demonstrate that Zfh1 is expressed in MPs, where it is required for their maintenance, and that its expression continues into adult-hood in a small subset of myogenic cells. If, as these data suggest, Zfh1 is important for sustaining a population of a persistent adult progenitors, there must be a mechanism that maintains Zfh1 expression in these cells while the remainder differentiate into functional flight muscles.

### *zfh1* enhancers conferring expression in MPs

To investigate whether the maintenance of Zfh1 expression in larval and adult MPs could be attributed to distinct enhancers, we screened *enhancer*-*Gal4* collections (Jenett, et al., 2012; Jory, et al., 2012; Manning, et al., 2012) to identify *zfh1* enhancers that were active in larval MPs. From the fifteen enhancers across the *zfh1* genomic locus that were tested, (Figure 2 and Figure supplement 3 A) three directed GFP expression in the Cut expressing MPs (Figure 2B-D). These all correlated with regions bound by the myogenic factor Twist in muscle progenitor related cells (Figure supplement 3 A and (Bernard, et al., 2010)). Enhancer 1 (Enh1; VT050105) conferred weak expression in scattered progenitors (Figure 2B). Enhancer 2 (Enh2; VT050115) was uniformly active in all MPs and also showed ectopic expression in some non-Cut expressing cells (Figure 2C). Finally, Enhancer 3 (Enh3; GMR35H09) conferred expression in several MPs with highest levels in a subset located in the posterior (Figure 2D). Enh3 encompasses a region that was previously shown to be bound by Su(H) in muscle progenitor related cells, hence may be regulated by Notch activity (Figure supplement 3A; (Bernard, et al., 2010; Krejci, et al., 2009). These results demonstrate that several enhancers contribute to *zfh1* expression in the MPs.

**Figure 2.**
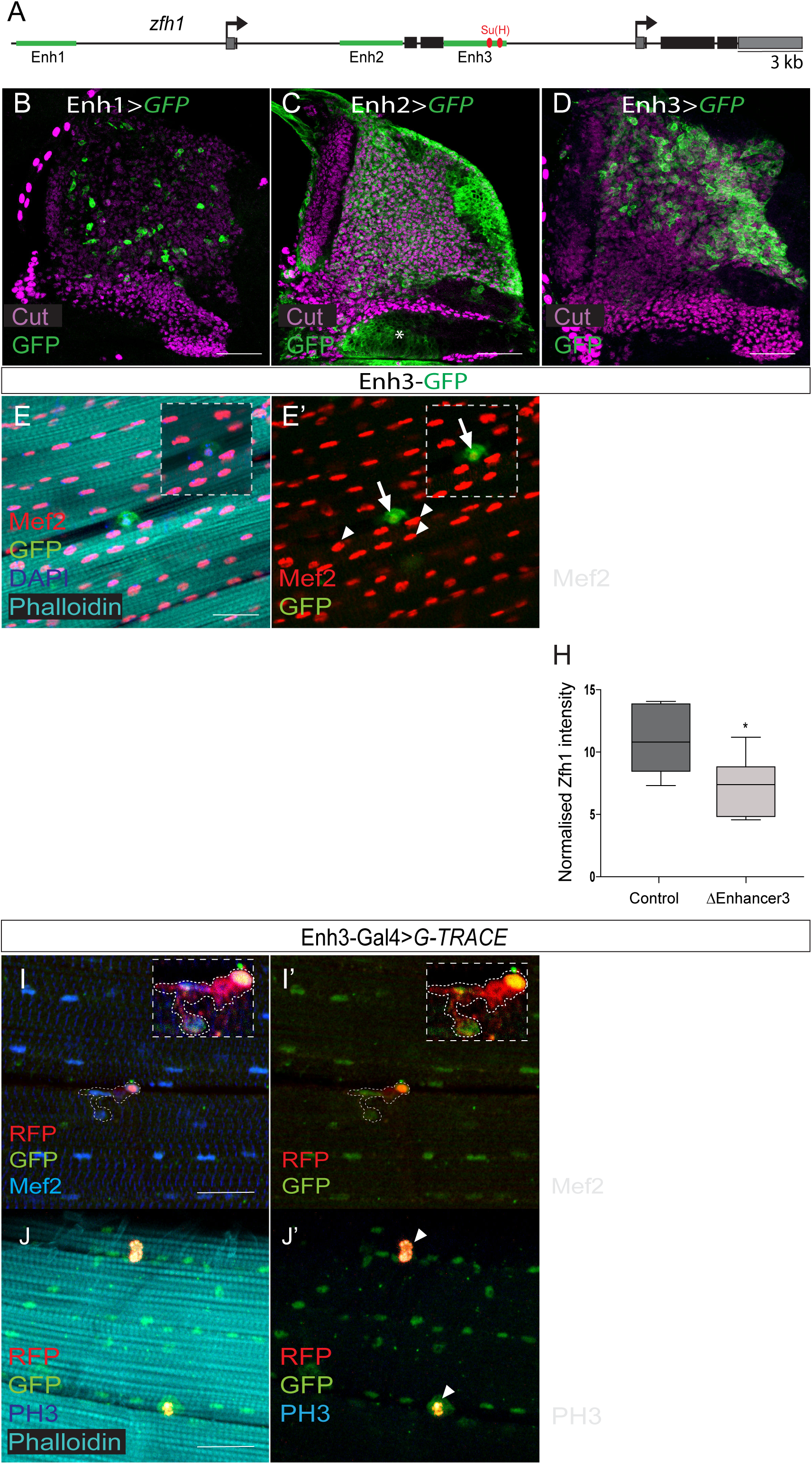
Regulation of *zfh1* in MPs and adult pMPs. **(A).** Schematic view of *zfh1* genomic region, *zfh1* regulatory enhancers are represented by green rectangles and arrows indicate transcription starts. Coding exons and untranslated regions are represented in black and grey boxes, respectively. **(B-D).** Three different zfh1 enhancers are active in the MPs (labelled with Cut, purple). *Enh1*-*Gal4* (VT050105, B) drives GFP (Green) in a subset of scattered MPs; *Enh2*-*Gal4* (VT050115, C) drives GFP throughout the MPs, and in some non-MP cells (Asterisk); *Enh3*-*Gal4* (GMR35H09, D) is highly expressed in a subset of MPs located in the posterior region of the notum. Scale bars: 50 μM. **(E-E”).** *Enh3*-*GFP* (Green) expression is retained in pMPs (characterised by low level of Mef2, red; arrows E’-E”) but not in differentiated muscle nuclei (high Mef2, red; arrowheads E’-E”). Phalloidin marks muscles and DAPI labels all nuclei (Blue). Insets: boxed regions magnified 12.5 X. Scale bars: 25 μM. **(F-H).** Zfh1(white) expression in MPs is significantly reduced in discs from *Enh3* deletion Δ*Enh3* (G, H) compared to controls (F, H). (^⋆^ *p*= 0.0379, n =13, Student t-test). Scale bars: 50 μM. **(I-J”).** Lineage tracing shows that adult pMPs contribute to muscles. Indirect flight muscles from adult flies where *Enh3*-*Gal4* drives expression of the G-Trace cassette; GFP (Green) indicates myoblasts that have expressed *Enh3*-*Gal4*, RFP (red) indicates myoblasts where Gal4 is still active, Mef2 labels muscle nuclei (Blue). Insets: boxed regions magnified 20 X. **(J-J”).** pMPs are mitoticaly active, indicated by anti-phosphH3 (Blue, arrowheads J’-J”).

To determine which enhancer(s) are also capable of conferring *zfh*-*1* expression in adult pMPs we assessed their activity in adult muscle preparations. Only Enh3 exhibited any activity in these cells (Figure 2E), where it recapitulated well Zfh1 protein expression (Figure 2 E-E”). Thus, Enh3-GFP was clearly detectable in scattered cells, which were closely apposed to the muscle fibers and contained low levels of Mef-2 (Figure 2E”), and was not expressed in the differentiated muscle nuclei (Figure 2). These results suggest that Enh3 is responsible for maintaining *zfh1* transcription in the adult pMPs.

If Enh3 is indeed necessary for expression of *zfh1* in MPs and pMPs, its removal should curtail *zfh1* expression in those cells. To test this, Enh3 was deleted by Crispr/Cas9 genome editing (ΔEnh3; see material and methods). ΔEnh3 homozygous flies survived until early pupal stages allowing us to analyze the phenotype at larval stages. As predicted, ΔEnh3 MPs exhibited greatly reduced Zfh1 protein expression (Figure 3F-H) that correlated with decreased *zfh1* mRNA levels (Figure supplement 3B). Although striking, the effects of ΔEnh3 did not phenocopy those of depleting *zfh1* using RNAi, as no premature up-regulation of muscle differentiation markers (MHC, Tm) occurred in ΔEnh3 discs (data not shown). This is likely due to residual *zfh1* mRNA/protein (Figure 2G), brought about by the activity of other *zfh1* enhancers (e.g. Enh1 and Enh2, Figure 2B-C). Nevertheless, it is evident that Enh3 has a key role in directing *zfh1* expression in MPs.

**Figure 3.**
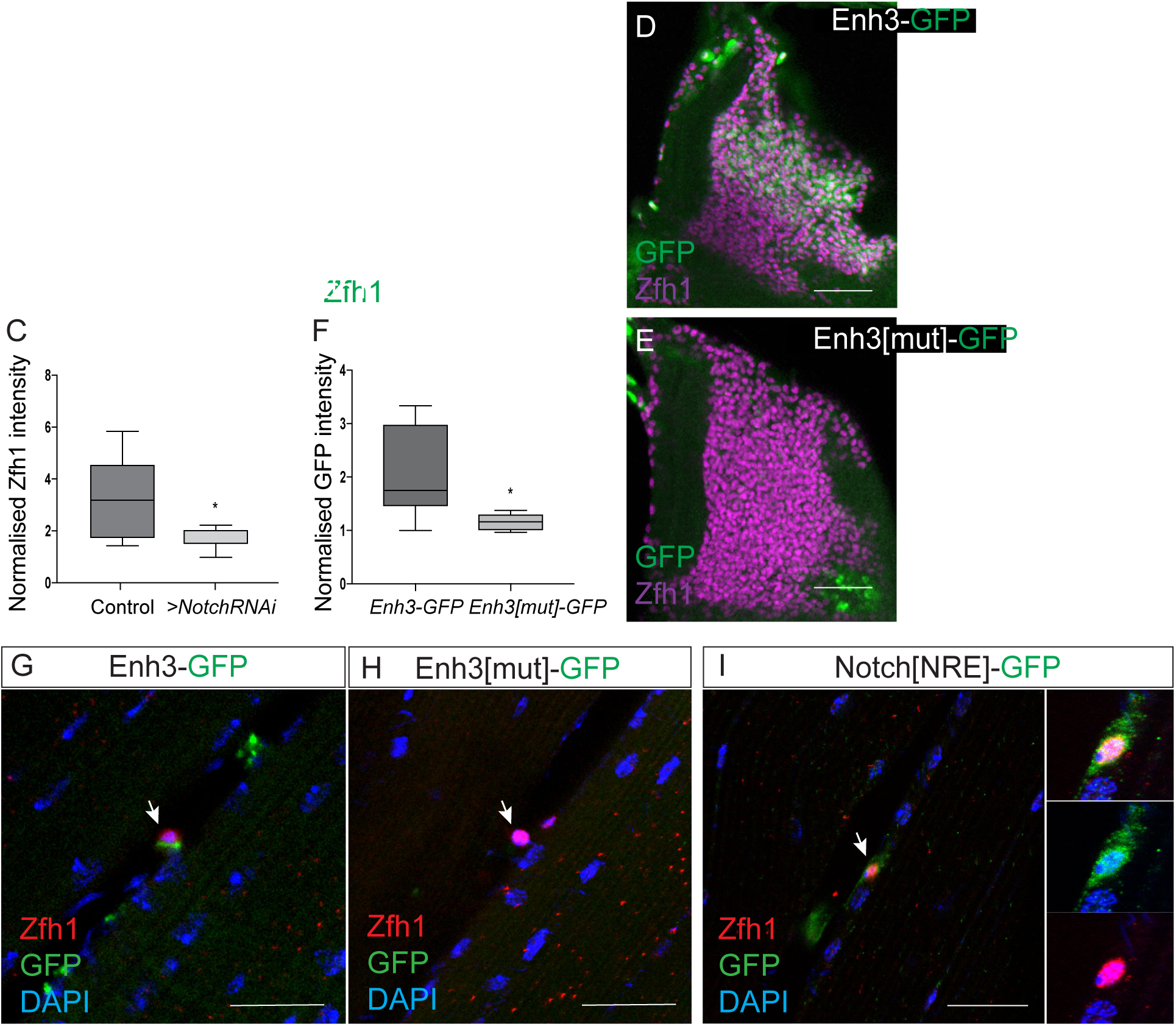
Notch directs Zfh1 expression in MPs and pMPs. **(A-C).** Zfh1 level (white) is significantly reduced when *Notch* is down regulated. Expression of Zfh1 in MPs (A) is severely reduced in the presence of *Notch RNAi* (B, *1151*-*Gal4*>*UAS*-*NotchRNAi*), Scale Bars: 50 μM. (C) Quantification of Zfh1 expression levels (^⋆^ *p* < 0.05, n= 12). **(D-F).** Enh3 (D, *Enh3*-*GFP*, green) expression in MPs (purple, Zfh1) is abolished when Su(H) motifs are mutated (E, *Enh3[mut]*-*GFP*). Scale bars: 50 μM. (F) Intensity of expression from *Enh3* and *Enh3[mut]* was significantly diffirent (^⋆^ *p*= 0.022, n=14, Student t-test). **(G-H).** Enh3 (G, *Enh3*-*GFP*, green) expression in adult pMPs (red, Zfh1) is abolished when Su(H) motifs are mutated (H, *Enh3[mut]*-*GFP*, green), DAPI (blue) reveals all nuclei. **(I).** *Notch[NRE]*-*GFP* (green) is co-expressed with Zfh1 (red) in the pMPs associated with the indirect flight muscle; DAPI (Blue) detects all nuclei. (N=12 pMPs). In G-I anti-P1 was included to label immune cells and exclude them from the analysis. Scale bars: 25 μM.

### Adult Zfh1+ve MP cells contribute to flight muscles

By recapitulating Zfh1 in adult MPs, Enh3 provides a powerful tool to investigate whether the persistent MPs are analogous to muscle satellite cells, which are able to divide and produce committed post-mitotic myogenic cells that participate in muscle growth and regeneration. To address this we used a genetic G-trace method, which involves two UAS reporters, an RFP reporter that directly monitors the current activity of the Gal4 and a GFP reporter that records the history of its expression to reveal the lineage (Evans, et al., 2009). When *Enh3*-*Gal4* was combined with the G-trace cassette RFP expression was present in the muscle-associated pMPs, which have low Mef2 expression (Figure 2I-I”). Strikingly, most of the muscle nuclei expressed GFP (Figure 2I’-I”) suggesting that they are derived from ancestral Enh3 expressing cells (Figure 2I’). Furthermore, close examination of the Enh3 driven RFP expression showed that it often persisted in two nearby muscle nuclei. This suggests that these nuclei are recent progeny of Enh3 expressing cells, indicating that these cells have kept the ability to divide that is characteristic of a satellite-like stem cell population (Figure 2). To further substantiate this conclusion, we verified that adult Zfh1+ve cells were actively dividing cells, using the mitotic marker phosphohistone-3 (PH3) staining. Many examples of Zfh1+ve cells co-stained for PH3 (Figure 2J-J”) indicating that these adult cells are dividing cells. Taken together the results argue that the adult Zfh1+ve cells resemble mammalian satellite cells, retaining the capacity to divide and provide progeny that maintain the adult flight muscles.

### Notch directly regulates *zfh1* expression in muscle progenitors and adults pMPs

As mentioned above, *zfh1* is regulated by Notch activity in *Drosophila* DmD8, MP-related, cells (Krejci, et al., 2009), where Enh3 was bound by Su(H) (Figure 2A and Figure supplement 3). Furthermore, phenotypes from depletion of *zfh1* in MPs, were reminiscent of those elicited by loss of Notch (N) signaling (Figure 1 and (Krejci, et al., 2009)). Notch activity is therefore a candidate to maintain Zfh1 expression in the adult pMPs, thereby preventing their premature differentiation. As a first step to test whether Notch activity contributes to *zfh1* expression, we depleted *Notch* in muscle progenitors by driving Notch RNAi expression with *1151*-*Gal4* (Figure 3A-B and C). Under these conditions, Zfh1 expression was significantly reduced consistent with Notch being required for *zfh1* expression in MPs. Second, the consequences of perturbing Notch regulation by mutating the Su(H) binding motifs in Enh3 were analyzed. Two potential Su(H) binding sites are present in Enh3 and both are highly conserved across species (Figure 2A). Mutation of both motifs, *Enh3[mut]*, resulted in a dramatic decrease of the enhancer activity in the MPs (Figure 3D-E and F). This supports the hypothesis that Notch directly controls *zfh1* expression in MPs by regulating activity of Enh3.

Since Enh3 activity persists in the adult MPs (Figure 2), we next analyzed whether mutating the Su(H) motifs impacted expression in these adult pMPs. Similar to larval stage MPs, *Enh3[mut]* had lost the ability to direct expression of GFP in the adult pMPs (Figure 3G-H). Thus, the Su(H) motifs are essential for Enh3 to be active in the adult pMPs. These data support the model that persistence of Zfh1 expression in adult MPs is likely due to Notch input, acting throughout Enh3.

The results imply that Notch should be expressed and active in the adult pMPs. To investigate this, we made use of a *Notch[NRE]*-*GFP* reporter line. *Notch[NRE]* is an enhancer from the *Notch* gene, and itself regulated by Notch activity, such that it is a read out both of Notch expression and of Notch activity (Simon, et al., 2014). Strikingly, *Notch[NRE]*-*GFP* reporter is consistently expressed in the Zfh1+ve adult pMPs, confirming that Notch is active in these cells (Figure 3I) but not in the differentiated muscles. It is also active in some surrounding non-muscles cells/tissues (Figure 3I). Together, the results show that *zfh1* expression in the adult pMPs requires Notch activity acting through Enh3.

### *zfh1* is silenced by the conserved microRNA *miR*-*8/miR*-*200* in MPs

Although transcriptional control of *zfh1* by Notch explains one aspect of its regulation, since all larval MPs express Zfh1 it remained unclear how a subset maintain this expression and escape from differentiation to give rise to the adult pMPs. A candidate to confer an additional level of regulation on *zfh1* expression is *miR*-*8/miR*-*200*, which is important for silencing *zfh1* (and its mammalian homologue ZEB1) in several contexts. The regulatory loop between *miR*-*8/miR*-*200* and *zfh1/ZEB* has been extensively studied in both *Drosophila* and vertebrates and is mediated by a *miR*-*8/miR*-*200* seed site located in the 3’ untranslated region (3’UTR) (Antonello, et al., 2015; Vallejo, et al., 2011; Brabletz and Brabletz, 2010).

To determine whether *miR*-*8* could down-regulate *zfh1* in muscle progenitors to promote their differentiation into muscles, we first examined the spatiotemporal expression pattern of *miR*-*8* at larval and adult stages (Figure 4) using *miR*-*8*-*Gal4*, whose activity reflects the expression of the endogenous *miR*-*8* promoter (Karres, et al., 2007). *miR*-*8*-*Gal4* was combined with a membrane GFP (for Progenitors, Figure 4A) or a nuclear GFP (for Adult muscles nuclei, Figure 4D) to visualize its expression. At larval stages, *miR*-*8* expression was inversely correlated with Zfh1 levels. Thus in most of the MPs, where Zfh1 expression was high, there was little or no expression from *miR*-*8*-*Gal4* (Figure 4 B-B’ and Figure 1A). Conversely, in the subset of MPs where Zfh1 is low (high Ct expressing DFM precursors) there were correspondingly higher levels of *miR*-*8*-*Gal4*, although the absolute levels were low compared to some of the surrounding tissues; Figure 4C-C’ and Figure 1 A). In contrast, adult muscles had uniform and high levels of *miR*-*8*-*Gal4*; all of the differentiated muscle nuclei exhibited nuclear GFP (Figure 4D-D’). Strikingly, at the same time, *miR*-*8*-*Gal4* expression was absent from the Zfh1+ve adult pMPs (Figure 4D-D’). These data show that, during muscle formation, *miR*-*8* and Zfh1 are expressed in a mutually exclusive manner; supporting the model that *miR*-*8* negatively regulates *zfh1.*

**Figure 4.**
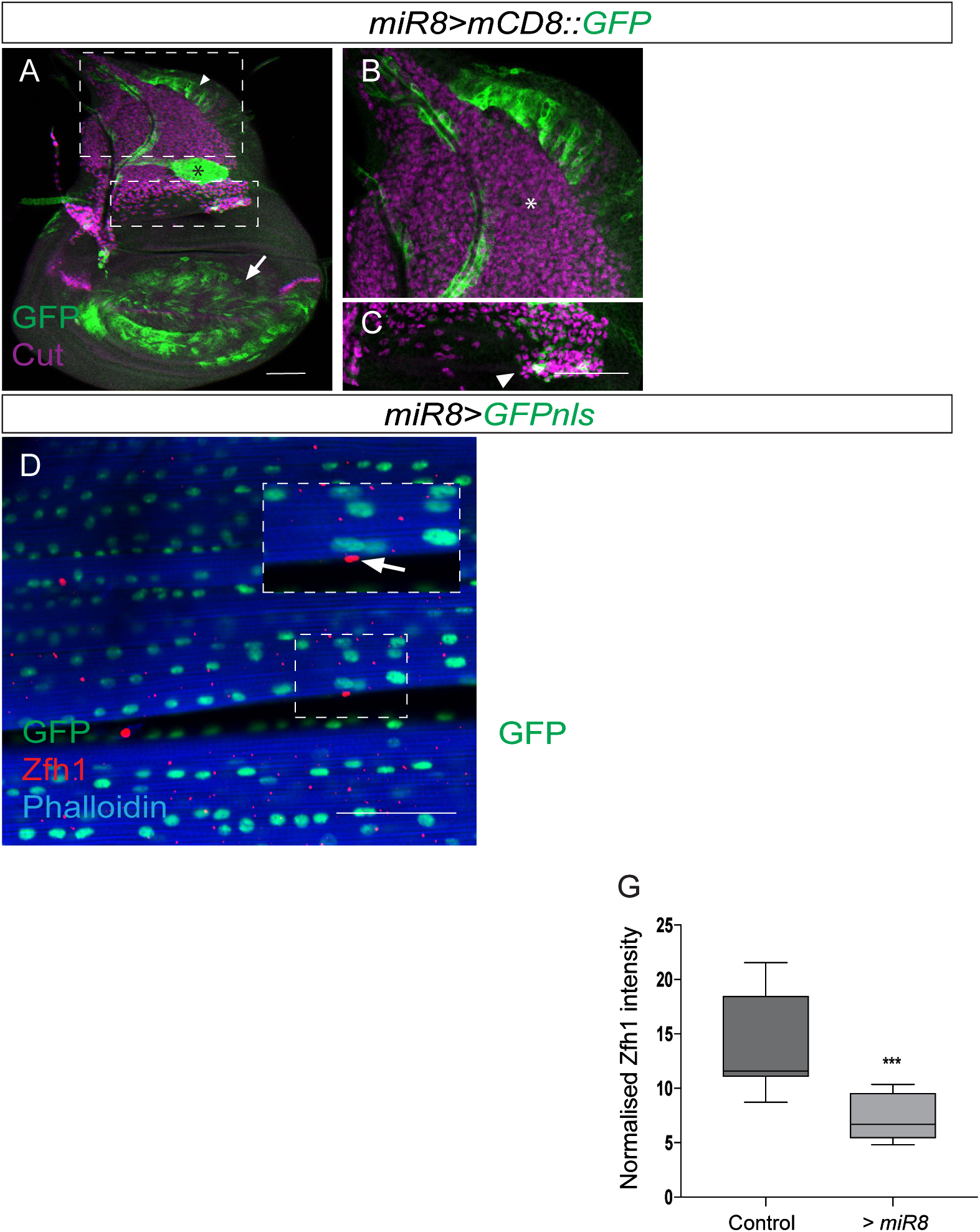
The conserved MicroRNA *miR*-*8/miR*-*200* antagonizes zfh1 to promote muscle differentiation. **(A-C).** Expression of *miR*-*8* is complementary to Zfh1; *miR*-*8* (green, *miR*-*8*-*Gal4*>*UAS*-*mCD8::GFP*) is not highly expressed in MPs (Cut, Purple) but is prevalent in the wing disc pouch (arrow), notum (arrowhead) and air sac (asterisk). Higher magnification shows that some *miR*-*8* expression can be detected in MPs where Zfh1 is normally low (arrowhead in C) but not in other MPs. Scale bars: 50 μM. **(D-D’)** *miR*-*8* (green; *miR*-*8*-*Gal4*>*UAS*-*nlsGFP*) expression is absent from Zfh1+ve (red) pMPs (arrows, D-D’) but is present at uniformly high levels in IFMs, (phalloidin, blue). Scale bars: 50 μM. **(E-G).** Effect of *miR*-*8* overexpression (*1151*-*Gal4*>*UAS*-*miR*-*8*) on Zfh1 (white) protein level in MPs. Scale Bars: 50 μM. (G) Zfh1 expression is significantly reduced by *miR*-*8* over-expression. (^⋆⋆⋆^ *p*= 0.0009, n=11, Student t-test).

To further address this regulation, we tested the impact of *miR*-*8* overexpression on Zfh1 protein levels in MPs at larval stage (Figure 4). Zfh1 protein levels were significantly diminished under these conditions, in agreement with *miR*-*8* regulating *zfh1* post-transcriptionally (Figure 4E-G). Although this manipulation was not sufficient to cause premature up-regulation of muscle differentiation markers, the few surviving adults all displayed a held-out wing phenotype, which is often associated with defective flight muscles (Vigoreaux, 2001).

Next we assessed whether *miR*-*8* activity/expression changes in response to Mef2 levels, a critical determinant of muscle differentiation, using a *miR*-*8* sensor (containing two *miR*-*8* binding sites in its 3’UTR (Kennell, et al., 2012). Expression of the *miR*-*8* sensor was specifically decreased when Mef2 was overexpressed in MPs, suggesting that *miR*-*8* expression responds to high level of Mef2 (Figure supplement 4). Together, these data argue that *miR*-*8* up-regulation during muscle differentiation blocks Zfh1 production to allow MPs to differentiate.

### An alternate short *zfh1* isoform is transcribed in adult MPs

To retain their undifferentiated state, the adult pMPs must evade *miR*-*8* regulation and maintain Zfh1 expression. The *zfh1* gene gives rise to three different mRNA isoforms; two long *zfh1* isoforms (*zfh1*-*long*; *zfh1*-*RE/RB*) and one short *zfh1* isoform (*zfh1*-*short*; *zfh1*-*RA*) (Figure 5A). Although long *zfh1* isoforms have two additional N-terminal zinc fingers, all three RNA-isoforms produce proteins containing the core zinc finger and homeodomains needed for Zfh1 DNA-binding activity (Figure supplement 5; (Postigo, et al., 1999)). Importantly, *zfh1*-*short* isoform has a shorter 3’UTR, which lacks the target site for *miR*-*8* (Figure 5A; (Antonello, et al., 2015), as well as differing in its transcription start site (TSS) (Figure 5A; Flybase FBgn0004606). As *zfh1*-*short* is predicted to escape *miR*-*8* mediated down-regulation, its expression will enable cells to retain high level of Zfh1 protein, explaining how the adult pMPs can escape differentiation. To test this prediction, we designed fluorescent probes specific for the *zfh1*-*long* and *zfh1*-*short* isoforms and used them for in situ hybridization (FISH) at larval and adult stages (Figure 5A). In the larval stage (L3), *zfh1*-*long isoforms were* present at uniformly high levels in the MPs (Figure 5B-B”’) whereas *zfh1*-*short* was expressed at much lower levels and only detected in a few MPs in each disc (Figure 5C-C”’).

**Figure 5.**
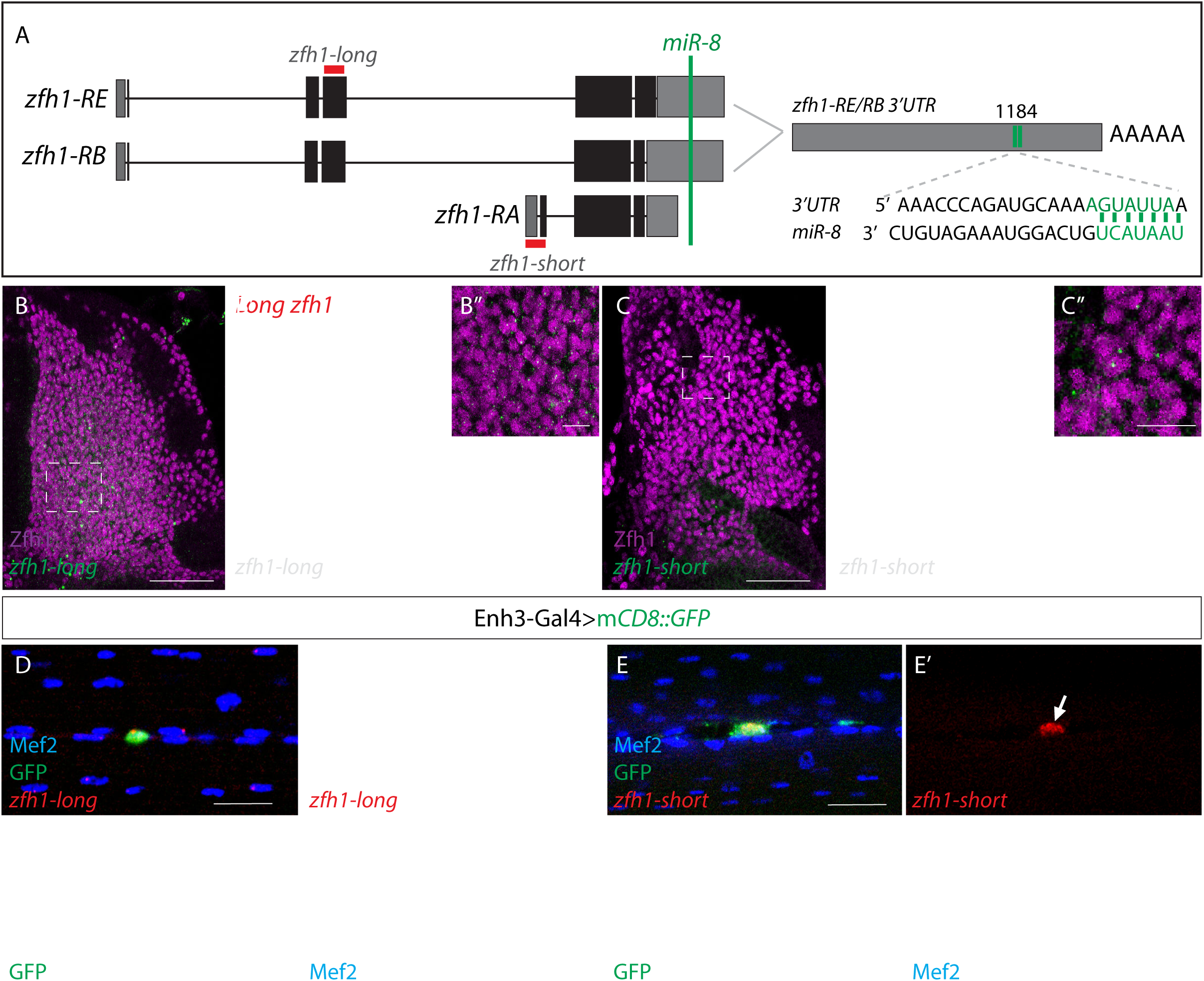
*zfh1* escapes *mir*-*8* regulation via a variable 3’UTR. **A.** Schematic representation of *zfh1* isoforms. *zfh1*-*short* (*zfh1*-*RA*) is initiated from a different transcription start site and has shorter 3’UTR that lack the target site for *miR*-*8* (green;(Antonello, et al., 2015)) present in *zfh1*-*long* isoforms (*zfh1*-*RB*, *zfh1*-*RE*); the position of the *miR*-*8* seed sites in *zfh1*-*long* 3’ UTR are depicted. Non-coding exons and coding exons are depicted by grey and black boxes respectively, red lines indicate the probes used for FISH experiments in B-E. **(B-C).** *zfh1*-*long* is present uniformly in MPs (B, green; B’, white) whereas *zfh1*-*short* is only detected in a few MPs (C, green; C’, white), detected by *in situ hybridisation* in wild type third instar wing discs stained for Zfh1 (Purple). Scale bars: 10 μM. (B”,B”’,C”C”’) Higher magnifications of boxed regions (Scale bars: 50 μM). **(D-E).** In adult IFMs *zfh1*-*long* is detected in the pMPs (arrow in D’) and in some differentiated nuclei located in their vicinity (arrowheads in D’) whereas *zfh1*-*short* is only present in pMPs (arrow in E’). *Enh3* expression (green, *Enh3*-*Gal4*>*UAS*-*mCD8GFP*) labels adult pMPs and Mef2 labels all muscle nuclei. Scale bars: 20 μM.

We then analyzed *zfh1* isoforms expression in adult muscles, where the adult pMPs were marked by *Enh3*-*Gal4*>*GFP* and low Mef2 expression (Figure 5D-E). In contrast to MPs in the larva, these adult pMPs expressed high levels of *zfh1*-*short* and much lower levels of *zfh1*-*long* (Figure 5D-E”’). *zfh1*-*short* was only present in the pMPs whereas dots of *zfh1*-*long* hybridization were also detected in some differentiated nuclei (with high level of Mef2) (Figure 5). These data indicate that *zfh1*-*long* isoforms are transcribed in both the progenitors and in muscle cells where *miR*-*8* regulation will prevent their translation, explaining the lack of protein. In contrast, *zfh1*-*short* is only expressed in a few larval MPs, and it is then detected at highest levels specifically in the adult pMPs. Since *zfh1*-*short* is not susceptible to regulation by *miR*-*8*, its specific transcription may therefore be determinant for maintaining high levels of Zfh1 in a subset of progenitors and enable them to escape differentiation.

### *zfh1* isoform transcription requires Notch activity in adult SCs

As expression of *zfh1*-*short* (*zfh1*-*RA*) in the adult MPs (Figure 5E) may be critical for them to retain their progenitor status, there must be mechanisms that ensure this isoform is appropriately transcribed. Notch is required for Zfh1 expression in MPs (Figure 3) and thus may contribute to this regulation. If this is the case, expression of a constitutively active Notch (NotchΔECD) should up regulate *zfh1*-*short* transcripts in the MPs at larval stages when its expression is normally low. In agreement, expression of active Notch in the progenitors (1151-Gal4>NotchΔECD) significantly increased the proportions of cells transcribing *zfh1*-*short* (Figure 6 A-B’ and *C*).

**Figure 6.**
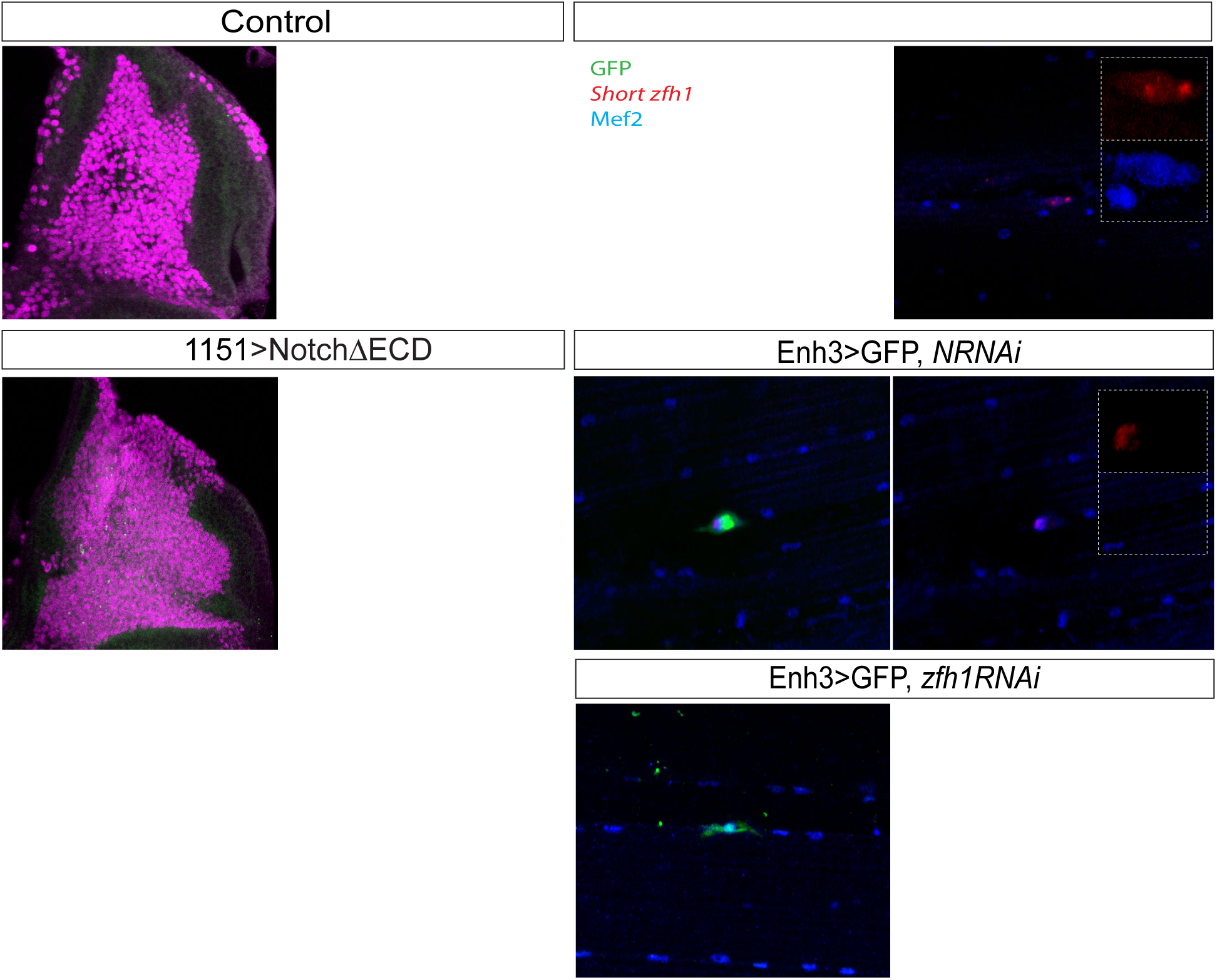

To address whether Notch is necessary for *zfh1*-*short* transcription in the adult pMPs, we specifically depleted Notch levels after eclosion (using *Enh3*-*Gal4* in combination with Gal80^ts^ to drive *Notch RNAi;* Figure 6 D-E’). Consistent with expression of *zfh1*-*short* being dependent on Notch activity, the levels of *zfh1-short* were significantly reduced in the adult pMPs when Notch was down-regulated after eclosion (Figure 6 D-E and G). Conversely, expression of *zfh1*-*long* was unaffected (Figure supplement 6). Notably, Mef2 accumulated to high levels in the adult pMPs under these conditions, suggesting that Notch activity helps prevent their differentiation (Figure 6 D’ and E’), most likely through its regulation of *zfh1*-*short.*

To investigate whether *zfh1* expression in adult MPs is required to prevent them differentiating, its expression was similarly downregulated in the pMPs, at 48 hours after eclosion. Targeted *zfh1* down-regulation in adults resulted in high levels of Mef2 being present in the adult MPs, suggesting that they had entered into the differentiation programme (Figure 6F-F’). Such forced differentiation of MPs would deplete the progenitor population and so should compromise muscle maintenance and repair. We therefore monitored the consequences of prolonged *zfh1* depletion on adult flies, assessing their phenotypes at ten days. Approximately 30% had a “held out wing” posture (n= 9/27) (Figure 7A-B), a phenotype often associated with flight muscle defects (Vigoreaux, 2001). The number of nuclei per muscle (DLM4) was also significantly reduced in the aged adults when *zfh1* was specifically depleted in the pMPs (Figure 7D-E). As no “held out wing” phenotype or muscle defects were observed in adult flies within 24 hours of knock-down, the phenotypes at 10 days are due a defect in the homeostasis of the adult flight muscles, consistent with Zfh1+ve cells contributing to their maintenance in a manner similar to the mammalian satellite cells.

**Figure 7.**
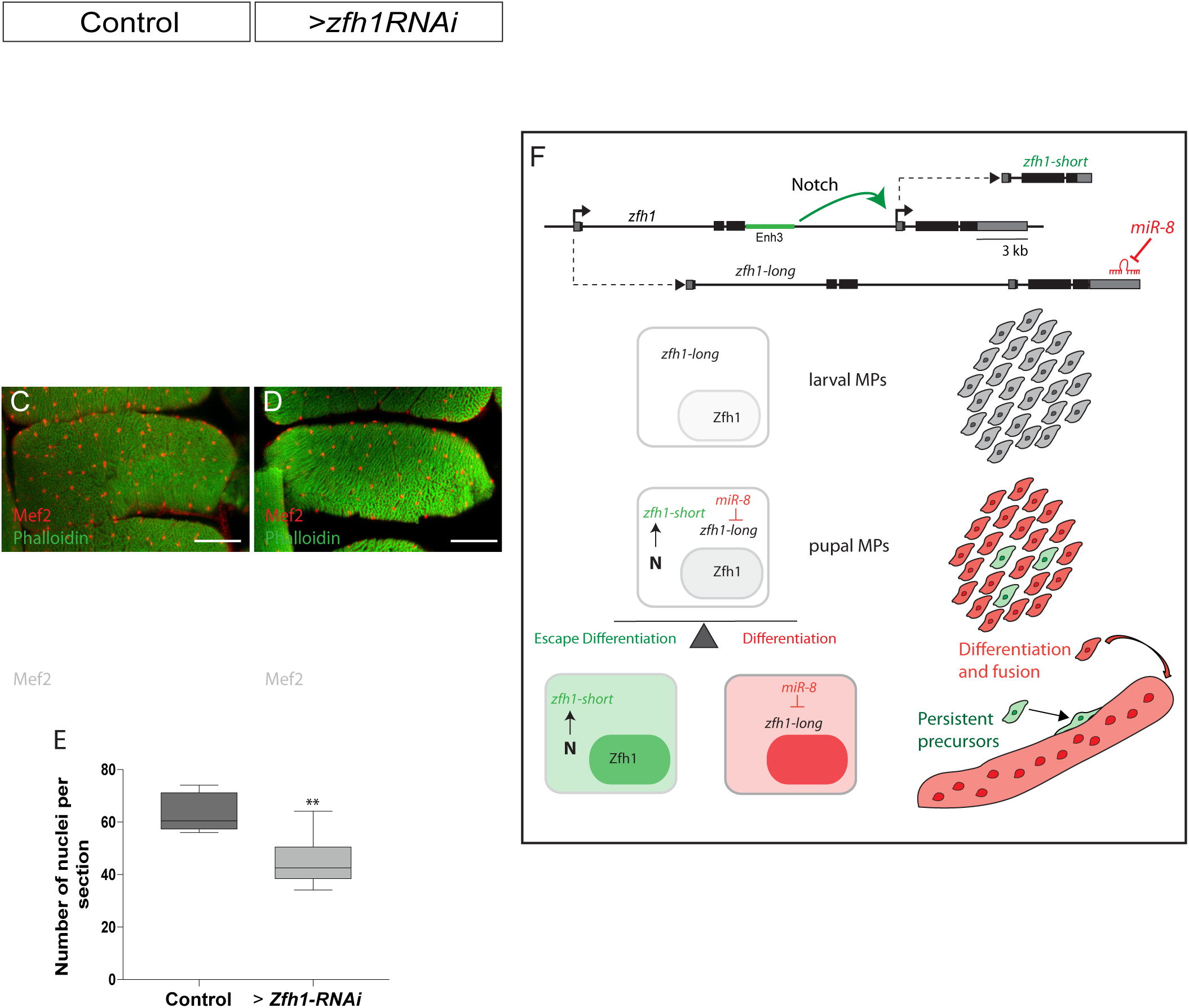
Zfh1 expression in pMPs is important to maintain muscle homeostasis. **(A-B).** Prolonged *zfh1* depletion in pMPs leads to a “held out” wings posture; dorsal view of (A) control (*Enh3*-*Gal4*>*UAS*-*wRNAi*; *tubGal80ts*) and (B) *zfh1* depletion (*Enh3*-*Gal4*>*UAS*-*zfh1RNAi*; *tubGal80ts*) adult flies. **(C-E).** Fewer Mef2+ve nuclei are present in muscles when zfh1 is depleted from pMPs. Transverse sections of DLM4 muscle stained with Phalloidin (green) and Mef2 (red, white) from the indicated genotypes. Scale bars: 50 μM (E) The number of nuclei per section in the indicated conditions was significantly different. (^⋆⋆^ *p* = 0.0013, n=10, Student t-test). **(F).** Model summarizing the role of alternate *zfh1* isoforms in the maintenance of adult pMPs. *zfh1*-*long* (Grey) is expressed in all MPs at larval stage. Silencing of *zfh1*-*long* by *mirR-8* (Red) facilitates the MPs differentiation. *zfh1*-*short* (Green) transcription is driven and maintained in pMPs by a Notch responsive element (Enh3, Green rectangle). Because *zfh1*-*short* is invisible to *miR*-*8*, Zfh1 protein is maintained in pMPs, enabling them to escape differentiation and persist as MPs in the adult.

## DISCUSSION

A key property of adult stem cells is their ability to remain in a quiescent state for a prolonged period of time (Li and Clevers, 2010). Investigating the maintenance of Drosophila MPs we have uncovered an important new regulatory logic, in which a switch in RNA isoforms enables a sub-population of cells to escape miRNA regulation and so avoid the differentiation programme. At the same time, analyzing expression of a pivotal player in this regulatory loop, *zfh1*, has revealed a population of persistent progenitors associated with adult muscles in Drosophila, that appear analogous to mammalian satellite cells (Figure 7F).

### Zfh1 marks a population of satellite-like cells in adult Drosophila

Until now, the fly has been thought to lack a persistent muscle stem cell population, leading to speculation about how its muscles could withstand the wear and tear of its active lifestyle. By analyzing the expression and function of Zfh-1, we identified a population of muscle-associated cells in the adult that retain progenitor-like properties, marked by Zfh-1 expression. Expression in the persistent adult “satellite-like” cells is dependent on one specific Zfh1 enhancer, which is directly regulated by Notch. Activity of Notch is important for maintaining Zfh1 expression and hence is required to sustain the progenitor status of these cells, similar to the situation in mammalian satellite cells, which require Notch activity for their maintenance (Mourikis and Tajbakhsh, 2014; Bjornson, et al., 2012).

Using lineage-tracing method we showed that adult Zfh1+ve cells, in normal conditions, provide new myoblasts to the fibers. Furthermore, conditional down regulation of *zfh1* led adult pMPs to enter differentiation and resulted in flight defects, evident by a held out wing posture. These results demonstrate that Zfh-1 is necessary to maintain these progenitors and that, similar to vertebrate satellite cells; the Zfh1+ve progenitor cells contribute to the adult muscles homeostasis. Thus the retention of a pool of progenitor cells may be critical to maintain the physiological function of all muscles in all organism types, as also highlighted by their identification in another arthropod *Parahyle* (Alwes, et al., 2016; Konstantinides and Averof, 2014). *Drosophila* notably differs because their satellite-like cells do not express the Pax3/Pax7 homologue (*gooseberry;* data not shown), considered a canonical marker in mammals and some other organisms (Chang and Rudnicki, 2014). Instead, Zfh1 appears to fulfill an analogous function and it will be interesting to discover how widespread this alternate Zfh1 pathway is for precursor maintenance. Notably, the loss of ZEB1 in mice accelerates the temporal expression muscle differentiation genes (eg. MHC) suggesting that there is indeed an evolutionary conserved function of Zfh1/ZEB in regulating the muscle differentiation process (Siles, et al., 2013). This lends further credence to the model that Zfh1 could have a fundamental role in preventing differentiation that may be harnessed in multiple contexts.

### Switching 3’ UTR to protect progenitors from differentiation

Another key feature of Zfh-1 regulation that is conserved between mammals and flies is its sensitivity to the miR-200/miR-8 family of miRNAs (Antonello, et al., 2015; Brabletz and Brabletz, 2010). This has major significance in many cancers, where loss of miR-200 results in elevated levels of ZEB1 promoting the expansion of cancer stem cells, and has led to a widely accepted model in which the downregulation of Zfh1 family is necessary to curb stem-ness (Brabletz and Brabletz, 2010). This fits with our observations, as we find that *miR*-*8* is upregulated during differentiation of the MPs and suppresses Zfh1 protein expression. Critically however, only some RNA isoforms, *zfh1*-*long*, contain seed sites necessary for *miR*-*8* regulation (Antonello, et al., 2015). The alternate, *zfh1*-*short*, isoform has a truncated 3’UTR that lacks the *miR*-*8* recognition sequences and will thus be insensitive to *miR*-*8* regulation. Significantly, this *zfh1*-*short* isoform is specifically expressed in MPs that persist into adulthood and hence can protect them from *miR*-*8* induced differentiation. Furthermore, it appears that Notch activity strongly promotes *zfh1* - *short* expression, explaining how this isoform is retained to sustain progenitors into adulthood.

Our study therefore provides a novel molecular logic explaining the maintenance of *Drosophila* satellite-like cells. This relies on the expression of *zfh1*-*short*, which, by being insensitive to *miR*-*8* regulation, can sustain Zfh1 protein production to protect pMPs from differentiation (Figure 7F). It also implies that Notch preferentially promotes the expression of a specific RNA isoform, most likely through the use of an alternate promoter in *zfh1.* Both of these concepts have widespread implications.

Alternate use of 3’UTRs, to escape miR regulation, is potentially an important mechanism to tune developmental decisions. Some tissues have a global tendency to favour certain isoform types, for example, distal polyadenylation sites are preferred in neuronal tissues (Zhang, et al., 2005). Furthermore, the occurrence of alternate 3’UTR RNA isoforms is widespread (>50% human genes generate alternate 3’UTR isoforms) and many conserved miR target sites are contained in such alternate 3’UTRs (Sandberg, et al., 2008; Tian, et al., 2005). Thus, similar isoform switching may underpin many instances of progenitor regulation and cell fate determination. Indeed an isoform switch appears to underlie variations in Pax3 expression levels between two different populations of muscle satellite cells in mice, where the use of alternative polyadenylation sites resulted in transcripts with shorter 3’UTRs that are resistant to regulation by *miR*-*206* (Boutet, et al., 2012). The selection of alternate 3’-UTRs could ensure that protein levels do not fall below a critical level (Yatsenko, et al., 2014), and in this way prevent differentiation from being triggered.

The switch in *zfh1* RNA isoforms is associated with Notch-dependent maintenance of the persistent adult MPs. Notably, *zfh1*-*short* is generated from an alternate promoter, as well as having a truncated 3’UTR, which may be one factor underlying this switch. Studies in yeasts demonstrate that looping occurs between promoters and polyadenylation sites, and that specific factors recruited at promoters can influence poly-A site selection (Lamas-Maceiras, et al., 2016; Tian and Manley, 2013). The levels and speed of transcription also appear to influence polyA site selection (Proudfoot, 2016; Tian and Manley, 2013; Pinto, et al., 2011). If Notch mediated activation via *Enh3* favors initiation at the *zfh1*-*short* promoter, this could in turn influence the selection of the proximal adenylation site to generate the truncated *miR*-*8* insensitive UTR. The concept that signaling can differentially regulate RNA sub-types has so far been little explored but our results suggest that is potentially of considerable significance. In future it will be important to investigate the extent that this mechanism is deployed in other contexts where signaling coordinates cell fate choices and stem cell maintenance.

## Materials and Methods

### Drosophila Genetics

All *Drosophila melanogaster* stocks were grown on standard medium at 25°C. The following stains were used: *w^118^* as wild type (wt), *UAS*-*white*-*RNAi* as control for *RNAi* experiments (BL #35573), *UAS*-*zfh1*-*RNAi* (VDRC: KK103205 and GD42856), *Mef2*-*Gal4* (Ranganayakulu, et al., 1996), *UAS*-*Mef2* (Cripps, et al., 2004), *UAS*-*GTRACE* (BL # 28281), *UAS*-*Notch*-*RNAi* (BL #7078), *Notch[NRE]*-*GFP* (Simon, et al., 2014), *UAS*-*Notch*Δ*ECD* (Chanet, et al., 2009; Fortini and Artavanis-Tsakonas, 1993; Rebay, et al., 1993), *miR8*-*Gal4* (Karres, et al., 2007), *UAS*-*miR8 (Vallejo*, *et al.*, *2011)*, *UAS*-*mCD8::GFP* (BL # 5137), *UAS*-*GFPnls* (BL # 65402), *UAS*-*Src::GFP* (Kaltschmidt, et al., 2000), *MHC*-*lacZ* (Hess et al., 1989), *1151*-*Gal4* (Anant, et al., 1998), *miR*-*8*-*sensor*-*EGFP* (Kennell, et al., 2012). RNAi experiments were conducted at 29°C and TubGal80^ts^ (McGuire, et al., 2003) was used to limit RNAi expression to a defined period of time. Enhancer-Gal4 lines described in Figure 2 and Figure supplement S3 are either from Janelia FlyLight (http://flweb.janelia.org) or Vienna Tiles Library (http://stockcenter.vdrc.at/control/main).

### Immunohistochemistry and *in situ* hybridization

Immunofluorescence stainings of wing discs were performed using standard techniques. Adult muscles were prepared and stained as described (Hunt and Demontis, 2013). The following primary antibody were used: Rabbit anti-Zfh1 (1:5000, a gift from Ruth Lehmann, New York, USA), Mouse anti-Cut (1:20, DSHB), Rabbit anti-β3-Tubulin (1:5000, a gift from Renate Renkawitz-Pohl, Marburg, Germany), Rat anti-Tropomyosin (1:1000, Abcam, ab50567), Goat anti-GFP (1:200, Abcam, ab6673), Rabbit anti-Ds-Red (1:25; Clontech, 6324496), Rabbit anti-Mef2 (1:200, a gift from Eileen Furlong, Heidelberg, Germany), Mouse anti-P1 (1:20, a gift from István Andó, Szeged, Hugary), Mouse anti-PH3 (1:100, Cell Signaling Technology, #9706), Mouse anti-β-Gal (1:1000, Promega, Z378A), Alexa-conjugated Phalloidin (1:200, Thermo fisher), Rat anti-Dcad2 (1:200, DSHB). *In situ* experiments were carried out according to Stellaris-protocols (https://www.biosearchtech.com/assets/bti_custom_stellaris_drosophila_protocol.pdf). Antibodies were included to the overnight hybridization step (together with the probes). *zfh1* probes were generated by Bioresearch Technologies. The sequence used for *zfh1*-*short* probe span 393bp of the first zfh1-RA exon, for *zfh1*-*long* probe, the sequence of the third exon (711bp) common to both zfh1-RB and zfh1-RE was used (see Figure 5).

### Construction of reporter lines and mutagenesis

For Enh3-GFP reporter line, the genomic region chr3R: 30774595..30778415 (Enh3/GMR35H09) according to Flybase genome release r6.03 was amplified using *yw* genomic DNA as template. Enh3 fragment was then cloned into the pGreenRabbit vector (Housden, et al., 2012). For *Enh3[mut]*-*GFP* line, two Su(H) biding sites were predicted within Enh3 sequence using Patser (Hertz and Stormo, 1999) and mutated by PCR based approach with primers overlapping the Su(H) sites to be mutated with the following sequence modifications: Su(H)1 AG<u>TG</u>G<u>GA</u>A to AG<u>GT</u>G<u>TG</u>A and Su(H) 2 T<u>TC</u>T<u>CA</u>CA to T<u>GT</u>T<u>TG</u>CA. Both constructs were inserted at position 68A4 on the third chromosome by injection into *nos*-*phiC31*-*NLS*; *attP2* embryos (Bischof, et al., 2007).

### CRISPR/Cas9 genome editing

CRISPR mediated deletion of Enh3 was performed according to (Port, et al., 2014) For generating guide RNAs, two protospacers were selected (sgRNA1 GCATTCCGCAGGTTTAGTCAC and sgRNA2 GCGATAACCCGGCGACCTCC) flanking 5’ and 3’ Enhancer-3 regions, (http://www.flyrnai.org/crispr/). The protospacers were cloned into the tandem guide RNA expression vector pCFD4 (Addgene # 49411) (http://www.crisprflydesign.org/wp-content/uploads/2014/06/Cloning-with-pCFD4.pdf). For the homology directed repair step, two homology arms were amplified using *yw* genomic DNA as template with the following primers (Homology arm1: Fwd. 5’ GCGCGAATTCGGGCTAAACGCCAGATAAGCG 3’ Rev. 5’ TTCCGCGGCCGCC ACTGGATTCCACGGCTTTTCG 3’– Homology arm 2: Fwd. 5’ GGTAGCTCTTCTTA TATAACCCGGCGACCTCCTCG 3’ Rev. 5’ GGTAGCTCTTCTGACCGGAC GAAAAACTAGCGACC) and cloned into the pDsRed-attP (Addgene # 51019) vector (http://flycrispr.molbio.wisc.edu/protocols/pHD-DsRed-attP). Both constructs were injected into *nos*-*Cas9* (BL # 54591) embryos. Flies mutant for Enhancer 3 were screened for the expression of the Ds-Red in the eyes and confirmed with sequencing of PCR fragment spanning the deletion.

### Microscopy and data analysis

Samples were imaged on Leica SP2 or TCS SP8 microscopes (CAIC, University of Cambridge) at 20X or 40X magnification and 1024/1024 pixel resolution. Images were processed with Image J and assembled with Adobe Illustrator. Quantification of fluorescence signal intensities was performed with Image J software. In each case the n refers to the number of individual specimens analyzed, which were from two or more independent experiments. For experiments to compare and measure expression levels, samples were prepared and analyzed in parallel, with identical conditions and the same laser parameters used for image acquisition. For each confocal stack a Sum slices projections was generated. Signal intensities were obtained by manually outlining the regions of interest, based on expression of markers, and measuring the average within each region. The values were then normalized to similar background measurements for each sample. In Figure 6 the number of transcriptional *zfh1*-*short* dots was counted manually with Image J and normalized to the total number of nuclei (Zfh1 staining), which was determined by a Matlab homemade script. Graphs and statistical analysis were performed with Prism 7 using unpaired t-test. Error bars indicate standard error of the mean.

### Quantitative RT PCR

30 Wing Imaginal discs from each genotype were dissected and RNA isolated using TRIzol (Life technologies). Quantitative PCR were performed as described (Krejci and Bray, 2007). Values were normalized to the level of *Rpl32.* The following primers were used. *Rpl32*, Fwd 5’-ATGCTAAGCTGTCGCACAAATG-3’ and Rev 5’-GTTCGATCCGTAACCGATGT-3’. *zfh1* Fwd 5’– GTTCAAGCACCACCTCAAGGAG-3’ and Rev 5’-CTTCTTGGAGGTCATGTGGGAGG-3’. (Product common to all three *zfh1* isoforms).

## Acknowledgements

We thank the Bloomington Stock Center, the VDRC Stock Center and the Developmental Studies Hybridoma Bank for *Drosophila* strains and antibodies. We acknowledge Ruth Lehmann, Renate Renkawitz-Pohl, Eileen Furlong and István Andó for antibodies. We thank Eva Zacharioudaki, Alain Vincent and Michalis Averof for critical reading of the manuscript and other members of SJB lab for valuable discussion. This work was funded by a programme grant from the MRC to SJB and by an EMBO Long Term Fellowship for HB.

---

**Figure supplement 1.**
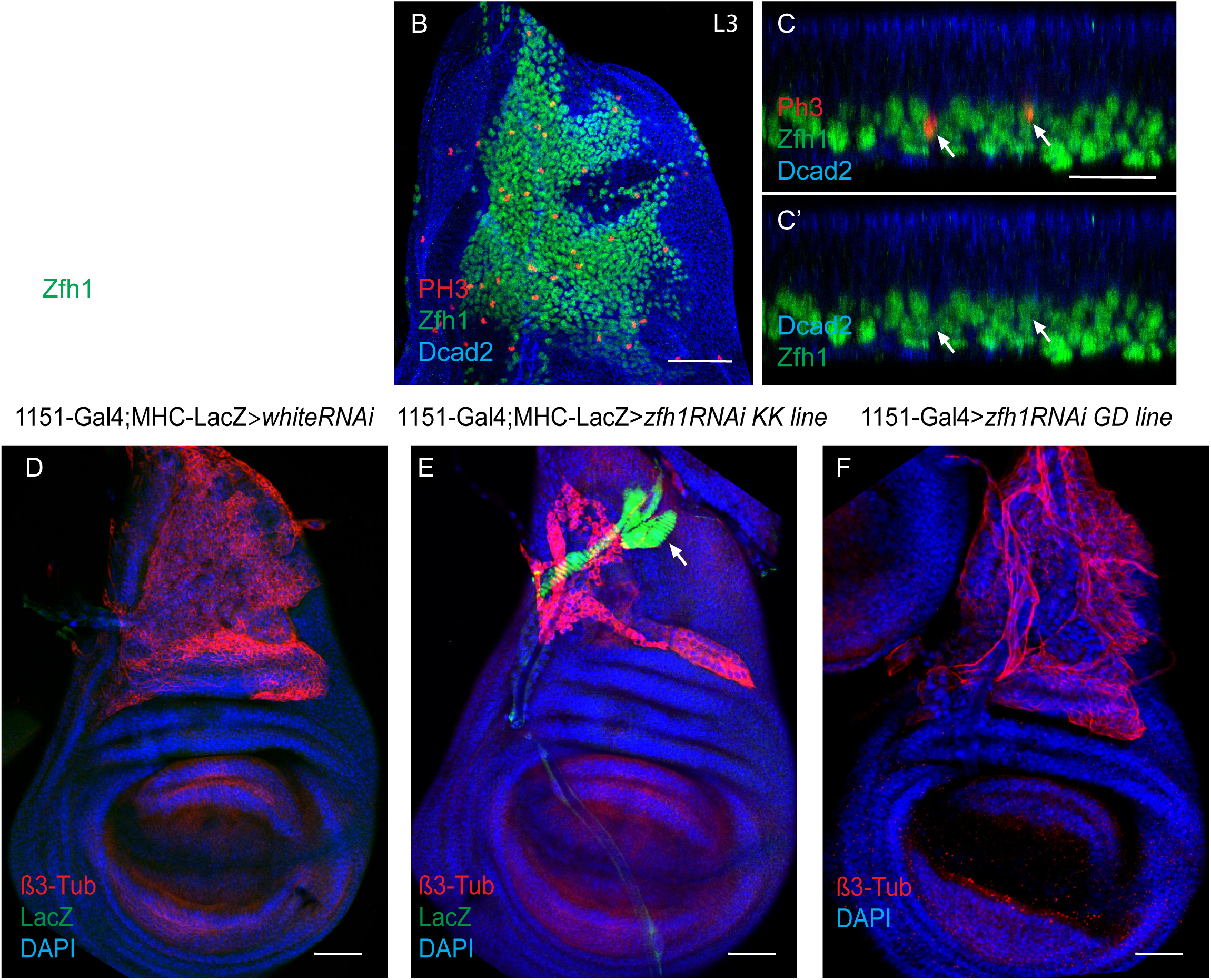
**A.** Zfh1 (Green) expression in MPs associated with early third instar wing discs showing that at this stage Zfh1 is uniformly expressed in all the MPs.Scale bar: 50 μM **B.** Third instar wing discs stained for Zfh1 (Green), PH3 (Red) and Dcad2 (Blue) revealing that some MPs are undertaking mitotic divisions. Scale bar: 50 μM. **(C-C’).** Optical section of image B showing active mitotic division of Zfh1 MPs in the most proximal layer to the disc epithelium (Arrows). Scale bar: 25 μM. **(D-F).** β3-Tubulin (β3-Tub, red), LacZ (Green) and DAPI (Blue) expression in control (D, *1151*-*Gal4*; *mhc*-*lacZ* > *UAS*-*white*-*RNAi*) and zfh1 down regulation (*1151*-*Gal4*>*UAS*-*zfh1*-*RNAi*) with two different RNAi transgenes, KK 103205 and GD42856, in E and F, respectively. Scale bars: 50 μM. **E.** Premature differentiation of MPs is observed using the KK103205 RNAi line comparing to the control (Arrow). **F.** Intermediate MPs differentiation phenotype is evident with the GD42856 RNAi transgene. MPs adopt an elongated morphology without expressing any muscle differentiation marker.

**Figure supplement 2.**
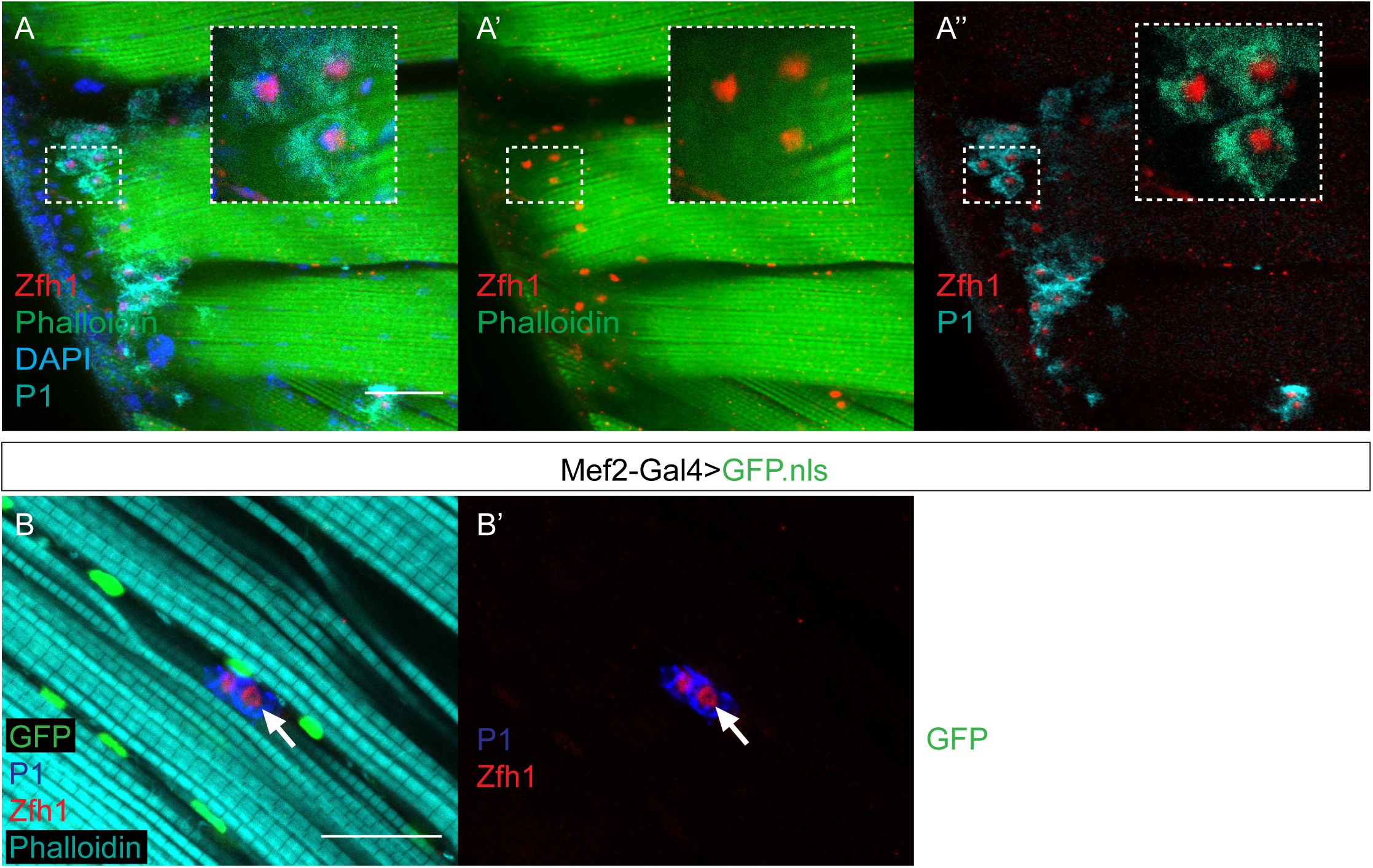
**A-A”.** Wild type indirect flight muscles stained for P1 (Cyan), Zfh1 (Red), Phalloidin (Green) and DAPI (Blue). P1 expression indicates the presence of phagocytic immune cells, which are also positive for Zfh1. Insets: boxed regions magnified 4X. Scale bar: 50 μM. **B-B”.** Indirect flight muscles from flies expressing *Mef2*-*Gal4*>*UAS*-*GFPnls* stained for GFP (Green), P1 (Blue), Zfh1 (Red) and Phalloidin (Cyan). The Zfh1 immune cells detected in the IFMs lack Mef2 expression (Arrows B-B”). Scale bar: 25 μM.

**Figure supplement 3.**
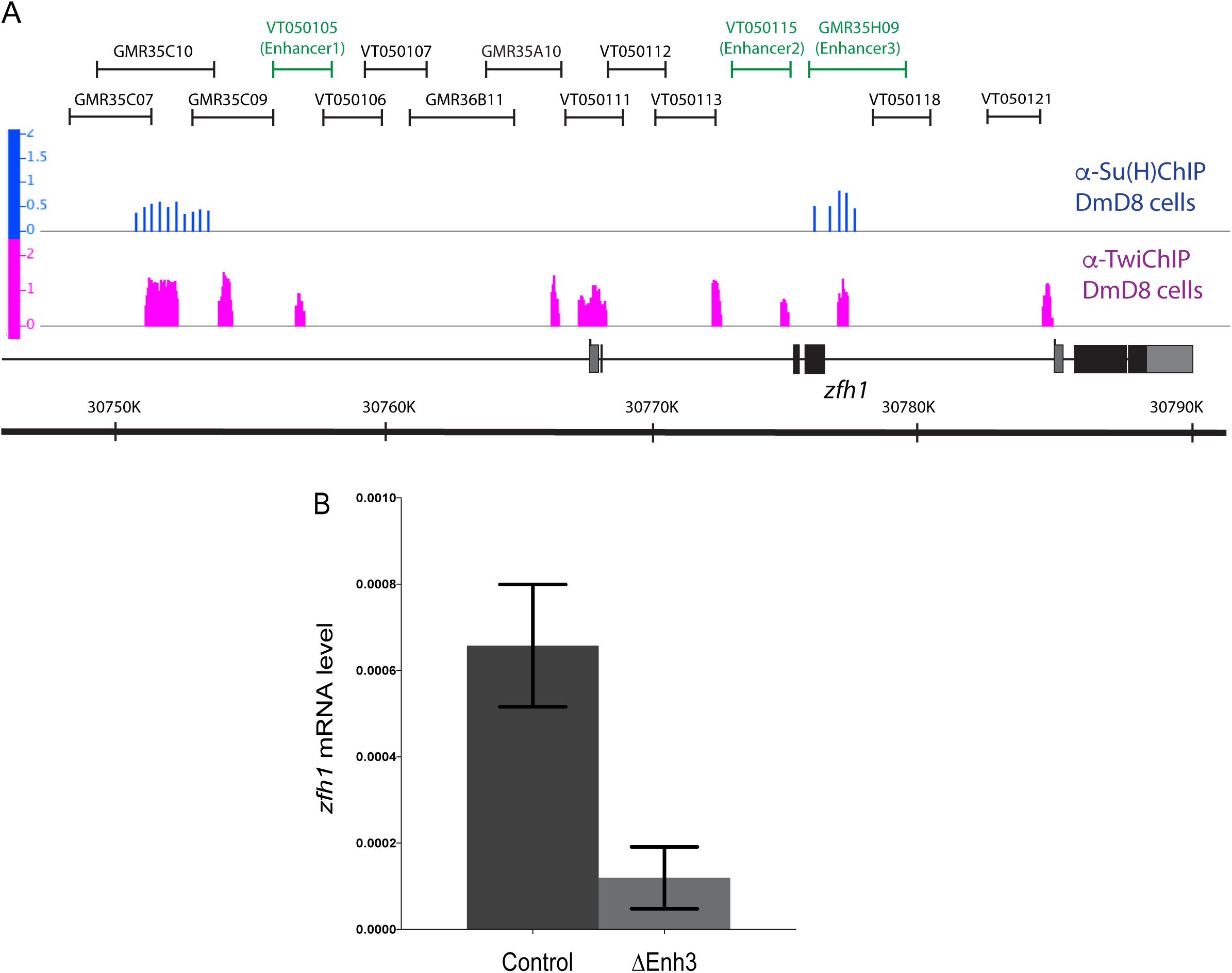
**A.** Identification of active *zfh1* enhancers in MPs and pMPs. Chromosome 3R: 30,765,926-30,789,202 showing the zfh1 genomic region with coding exons and untranslated regions represented in black and grey boxes, respectively. Arrows represent the transcription starts. Su(H) and Twi Chip enriched regions in DmD8 cells are represented in Blue and Magenta, respectively (Data from Bernard et al.,2010 (Chip Twi) and Krejci et al., 2009 (Chip Su(H)). Black lines represent all GMR and VT lines tested in our study. Green lines indicate the enhancers that are active in MPs *in vivo.* **B.** Effect of Enh3 Crisper deletion (ΔEnh3) on *zfh1* mRNA level in larval MPs as determined by quantitative RT-PCR. zfh1 mRNA level is significantly decreased by Enh3 deletion.

**Figure supplement 4.**
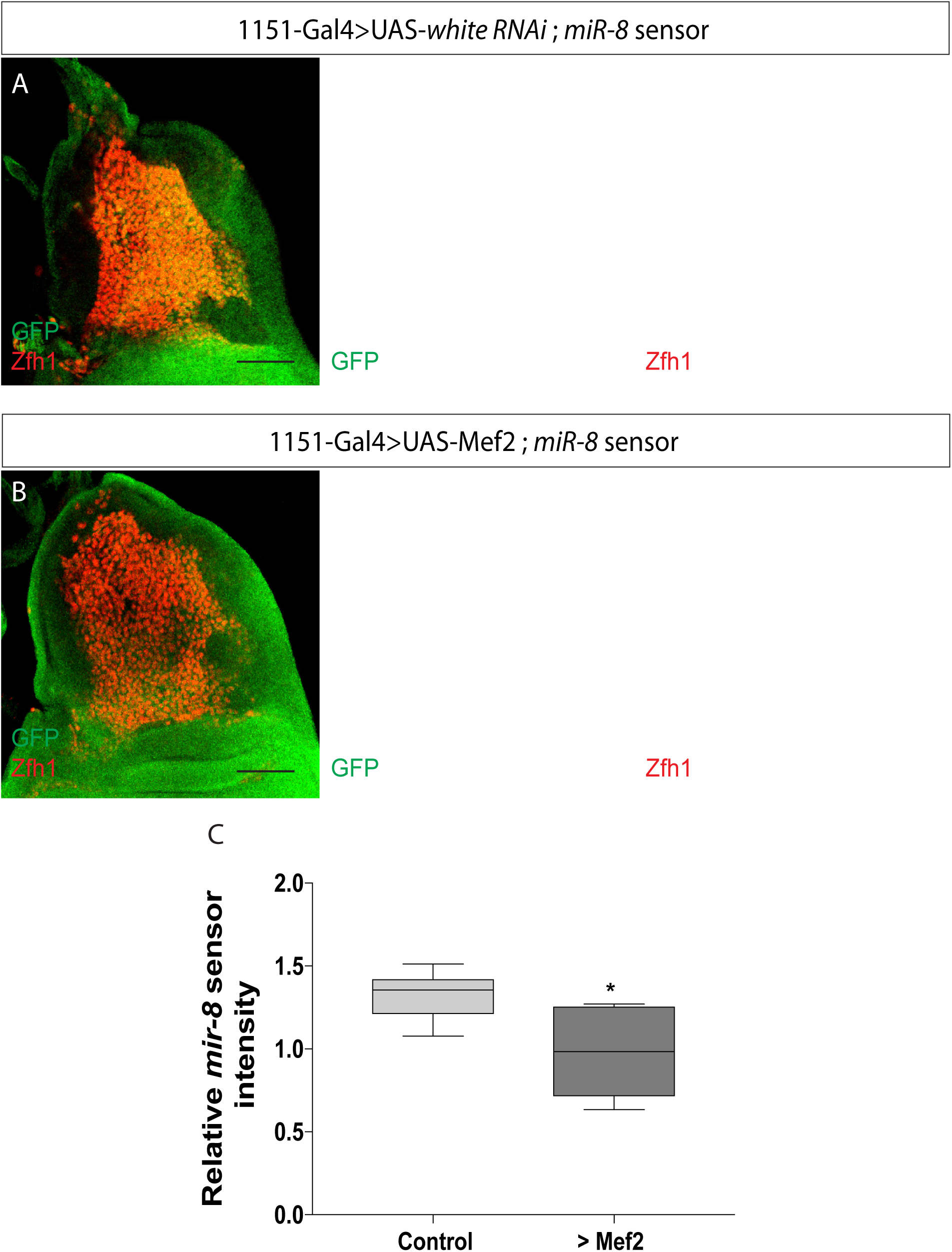
*miR*-*8* responds to high level of Mef2. *miR*-*8* activity was detected using *miR*-*8*-*EGFP* sensor (Kennel et al., 2012). **(A-B).** Effect of Mef2 overexpression (*1151*-*Gal4*>*UAS*-*Mef2*) on miR-8 sensor (Green) expression level in MPs. *miR*-*8* sensor expression in MPs is significantly reduced by Mef2 overexpression (Arrows in A’ and B’). Zfh1 expression (Red) was used to visualise the MPs. **(C).** Quantification of *miR*-*8* Sensor intensity in the indicated conditions. (^⋆^ *p* = 0.0164, n= 8). Scale bars = 50 μM.

**Figure supplement 5.**
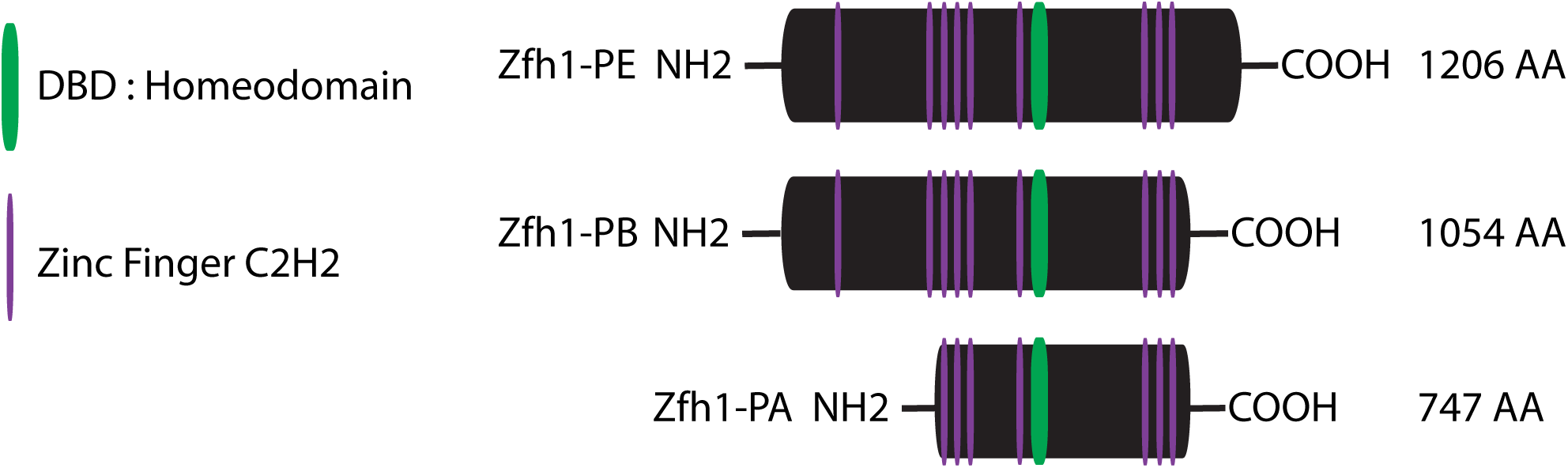
Domain structure of Drosophila Zfh1 protein isoforms E, B and A. Magenta bars indicate the predicted zinc fingers, Green bar indicates the predicted homeodomain. All three protein-isoforms contain the core zinc finger and homeodomains needed for Zfh1 DNA-binding activity. Isoforms E and B contain two additional N-terminal zinc fingers (Postigo, et al., 1999).

**Figure supplement 6.**
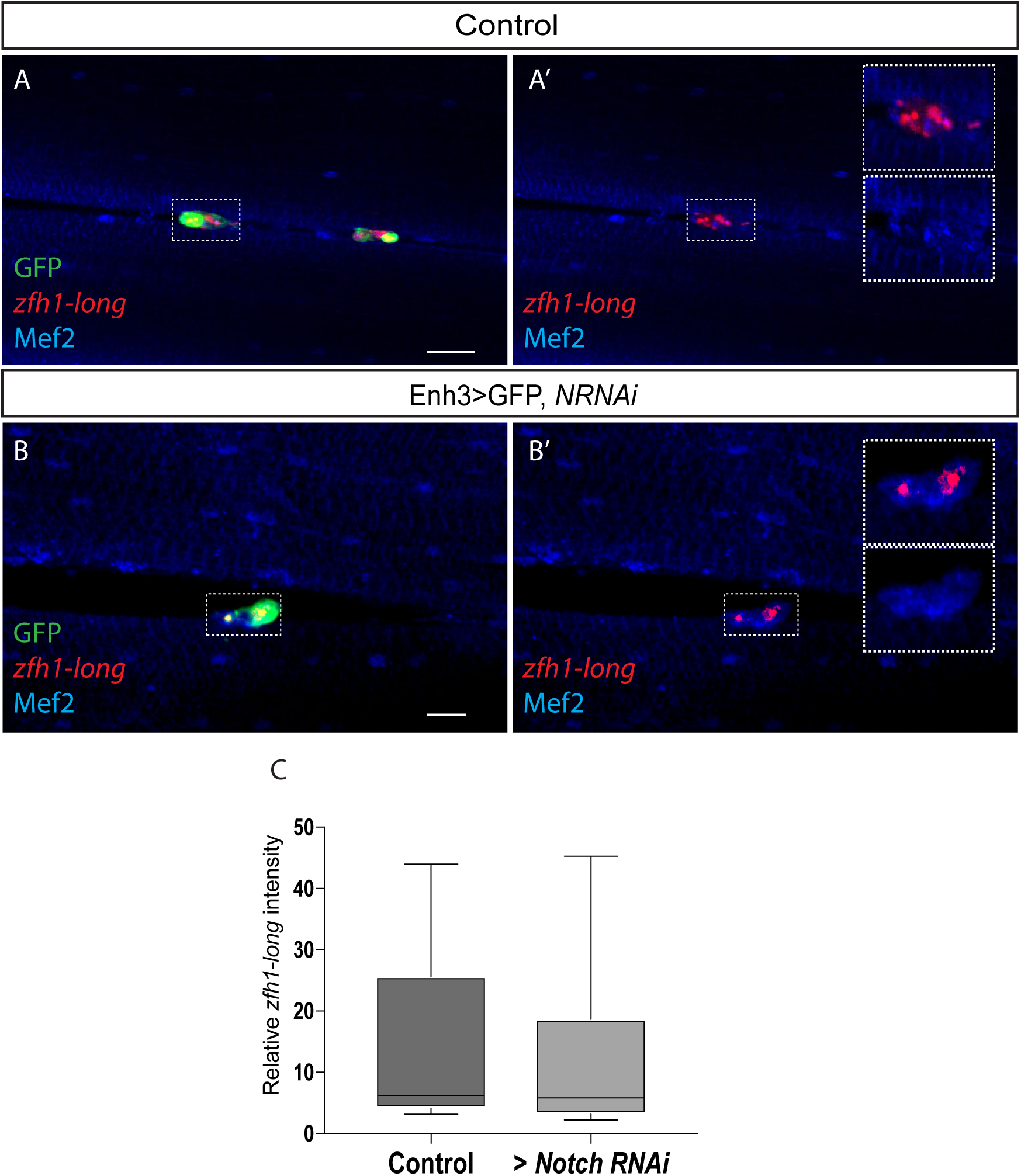
Notch activity is not necessary for *zfh1*-*long* (red) transcription (A’-B’ and C) in pMPs (green; *Enh3*-*Gal4*>*UAS*-*mCD8GFP*). *Enh3*-*Gal4* was used to drive expression of control RNAi (A-A’), *Notch* RNAi (B,B’) specifically in pMPs. Mef2 (Blue) marks all muscle and pMP nuclei. Scale bars: 25 μM. (C) Quantification of *zfh1*-*long* intensity in the indicated conditions, no significant difference was detected.

